# *ViReMaShiny*: An Interactive Application for Analysis of Viral Recombination Data

**DOI:** 10.1101/2022.04.06.487215

**Authors:** Jason Yeung, Andrew L Routh

## Abstract

Recombination is an essential driver of virus evolution and adaption, giving rise to new chimeric viruses, structural variants, sub-genomic RNAs, and Defective-RNAs. Next-Generation Sequencing of virus samples, either from experimental or clinical settings, has revealed a complex distribution of recombination events that contributes to the intrahost diversity. We and others have previously developed alignment tools to discover and map these diverse recombination events in NGS data. However, there is no standard for data visualization to contextualize events of interest and downstream analysis often requires bespoke coding. To address this, we present *ViReMaShiny*, a web-based application built using the R Shiny framework to allow interactive exploration and point-and-click visualization of viral recombination data provided in BED format generated by computational pipelines such as *ViReMa* (Viral-Recombination-Mapper).

## Introduction

Viruses exist as dynamic populations of diverse genomes (often referred to as intra-host diversity) which is maintained by the error-prone nature of viral replication (Lauring, et al., 2013). In addition to single-nucleotide variations (SNVs), viral recombination also contributes to the intrahost diversity through the production of structural variants (SVs), sub-genomic RNAs (sgmRNAs), Defective-RNAs (D-RNAs) and chimeric viruses that seed the emergence of novel virus strains (Simon-Loriere and Holmes, 2011). Viral recombination has contributed to the generation of notable variants in SARS-CoV-2, such as conserved insertions and deletions in the Spike region of the Alpha, Delta and Omicron variants of concern (VOCs). Consequently, recombination has an important influence on viral evolution both within single hosts and on ecological scales.

Improved high-throughput sequencing capabilities have enabled unprecedented characterization of the diversity of viral recombination events within virus populations. These technical advancements have led to a corresponding demand for bioinformatic software that can discover and map these events. *ViReMa* (Viral-Recombination-Mapper) (Routh and Johnson, 2014) is a python package enabling viral read mapping from Next-Generation Sequencing (NGS) data that was developed for this purpose. ViReMa has been used to characterize the full gamut of different recombination events in multiple studies, including sub-genomic mRNA production in coronaviruses (Gribble, et al., 2021), drug-resistance development in HIV samples (Wang, et al., 2022), the evolution of D-RNAs during serial virus passaging of Flock House virus in culture (Jaworski and Routh, 2017), demonstrated differences in D-RNA abundance between intra- and extra-cellular compartments of alphaviruses (Langsjoen, et al., 2020), and compared recombination events from experimental and clinical isolates of SARS-CoV-2 (Jaworski, et al., 2021).

Due to the combination of skills required, these studies are commonly collaborations between wet-lab experimentalists and bioinformaticians. The collaborative process can require multiple iterations of analyses to home in on data that are both valid and biologically interesting. Data exploration using easily accessible, GUI-based applications can improve turn-around times between iterations and allows experimentalists with limited coding experience to actively engage in analysis.

We present a R Shiny application, *ViReMaShiny*, that enables rapid visualization of viral recombination data from *ViReMa* or other applications that output recombination events using BED files. This application seeks to standardize representation of key features in viral recombination events such as their frequency and position relative to important genomic elements.

## Results

The ViReMaShiny application was created using the R Shiny framework and relies on the *ggplot2* and *circlize* (Gu, et al., 2014) packages for plotting. The Shiny framework provides interactivity, extensibility, and flexibility with local and web-hosted options available. Initial input of user files requires BED files, an output of the *ViReMa* (Sotcheff, et al., 2022) or other recombination mappers and are hosted locally. Either a single BED file or multiple BED files from multiple biological or experimental replicates can be uploaded. The BED files follow the standardized format as depicted in **Figure 1A**. Briefly, the genome reference, strand, and coordinates of the donor and acceptors sites of the recombination junction are provided in addition to the number of reads that map to each unique event. When reads are mapped using *ViReMa*, additional columns also provide the read coverage at each junction site, and the sequence of a fixed number of nucleotides both up- and down-stream of the donor and the acceptor sites. This latter information allows scrutiny of the sequence composition of each recombination junction. Once uploaded, users will be able to subset data visualized by file name and included reference sequences through drop-down menu selectors.

**Figure 1.**
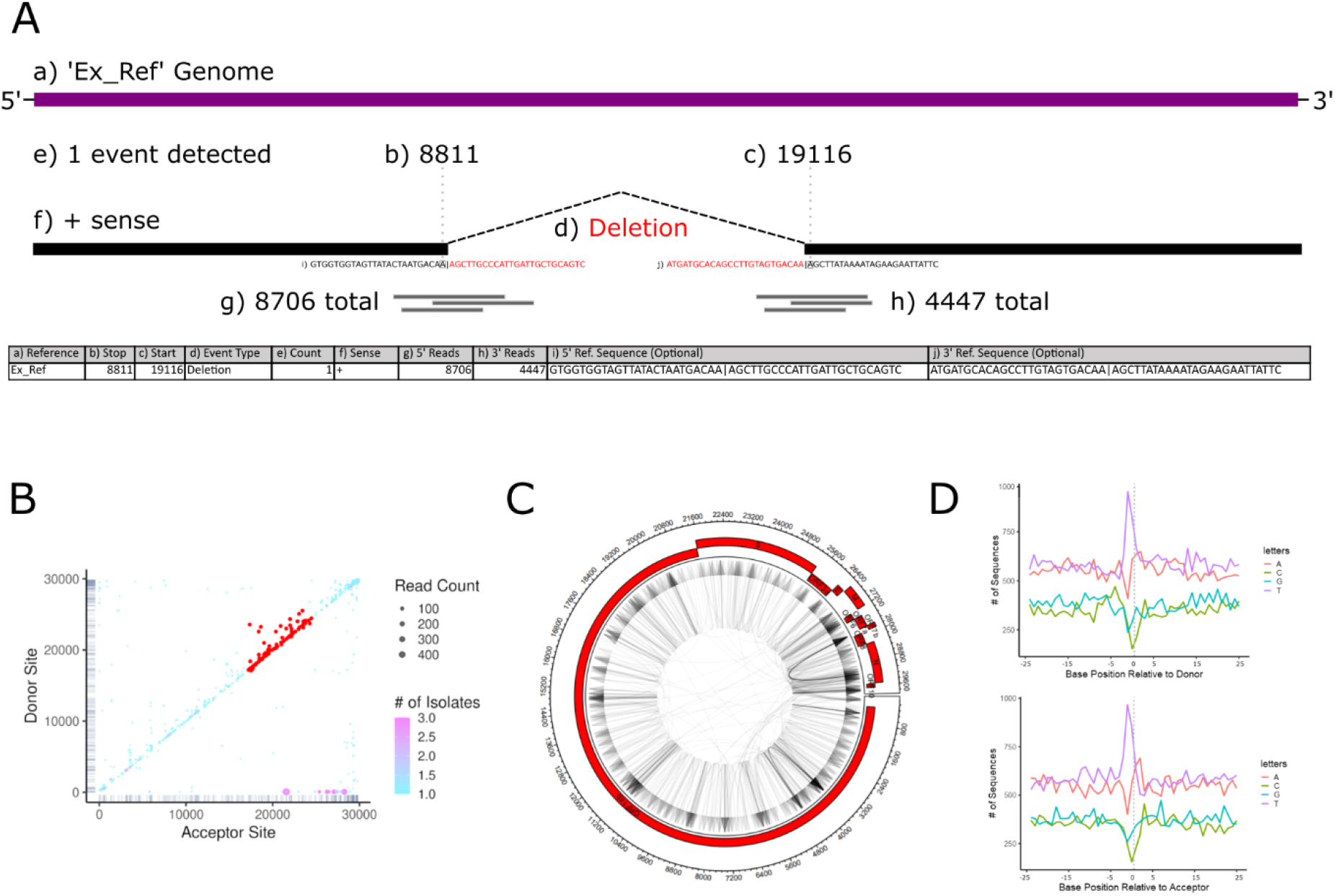
Example input data and plot outputs from *ViReMaShiny* using sample SARS-CoV-2 data from a previous study (Jaworski, et al., 2021). **A)** A table with an example recombination event in *ViReMa* output BED format. Subsection a) refers to the reference sequence of the event. b) and c) indicate the nucleotide base positions spanning the event. d) is the type of recombination event. e) is the number of reads detected with this event. f) is the sense or strandedness of the nucleic acid the event was detected on. g) and h) correspond to the number of reads spanning b) and c) respectively. i) and j) are reference-derived nucleotide sequences at the recombination junction, 25bp upstream and downstream of both b) and c). **B)** A scatterplot depicting recombination events by donor and acceptor site indexes. Read counts correspond to dot size while color encodes the number of isolates the event appears in. A subset of interest is highlighted in red. **C)** An example Circos plot depicting all recombination events. The plot includes annotations for the NCBI RefSeq NC_045512.2 SARS-CoV-2 genome. **D)** Nucleotide bias at donor and acceptor sites for all recombination events.

Interactivity is centered around two heatmaps depicting recombination events by donor (y-axis) and acceptor (x-axis) sites of each unique recombination junction. If multiple BED files representing multiple unique samples are uploaded, then a color-bar indicates the frequency with which specific recombination junctions are seen in multiple samples. We present an example of a recombination heatmap from samples of SARS-CoV-2 RNA obtained from three nasopharyngeal swabs from a previous study (Jaworski, et al., 2021) (**Figure 1B, SData 1**). This visualization quickly summarizes notable features of recombination events including favored acceptor or donor sites, evidence of recombination hotspots and the frequency or abundance of similar events within a dataset. For example, the conserved sgmRNAs of SARS-CoV-2 are visible as purple spots in the lower right portion of the heatmap and abundant insertions and deletions are observed in the upper right end of the heatmap, representing diverse RNA recombination near the 3’UTR of SARS-CoV-2. Frequent small InDels are represented by the numerous points close to the x=y axis. This approach has been extensively used to visualize recombination events in a range of viruses including Nodaviruses (Routh, et al., 2012), alphaviruses (Langsjoen, et al., 2020) and coronaviruses (Gribble, et al., 2021). Scrubbing the scatterplots generates a filterable table, allowing users to identify events with specific features. A text box above this table alternatively allows for filter-expressions in R syntax to be applied to data based on each of the parameters in the BED file(s). This allows users to sample specific events with desired features, such as (for example) only small InDels or only highly abundant events. Events in the table can be highlighted on the scatterplot using a toggle-able button or by clicking on a row in the sequence table.

Users can view summary statistics and plots under the “Overview” tab. A table summarizes total and unique events for each sample. If certain columns are included in the input files, frequency plots reveal any enrichment or depletion of specific nucleotides proximal and distal to donor and acceptor sites. For example, in SARS-CoV-2, recombination sites are most frequently flanked by U-rich tracks (**Figure 1D**). Manhattan plots show broad patterns in deleted and duplicated segments, as has previously been used to identify conserved functional motifs in the RNA genome of Flock House virus (Routh and Johnson, 2014). Finally, a *Circos* plot (Krzywinski, et al., 2009) depicts directional recombination events relative to user-provided annotations (Figure 1C). Numerous visual options can be adjusted through sliders. Annotations can either be added manually through an editable table or included as a BED file. Export of all table data and plots are available in high-fidelity file formats such as TIFF and PDF.

Ultimately, scientific questions such as most common recombination event, nucleotide usage proximal and distal to recombination sites, and samples with most unique events can be answered within minutes of file upload. Vignettes included in documentation demonstrate how to use the application for these purposes. Associated documentation also includes tutorials for analysis of *ViReMa* output data in R. Code for the shiny application is available for local use and extension at https://github.com/routhlab/ViReMaShiny.

## Conclusions

*ViReMaShiny* standardizes outputs and improves the approachability of exploratory viral recombination analysis. This application is built on the outputs of the *ViReMa* python script, allowing for intuitive investigation of data with no coding requirement. We plan on expanding support for analysis in the R environment as well as providing options to visualize recombination between multiple genes of multi-partite viruses such as influenza virus (Alnaji, et al., 2021). The application is hosted at https://routhlab.shinyapps.io/ViReMaShiny/ with associated documentation at https://jayeung12.github.io/. Code is available at https://github.com/routhlab/ViReMaShiny.

## Supporting information

SData 1

## Acknowledgements

We would like to thank Dr. Fadi Alnaji (University of Illinois at Urbana-Champaign) for discussions and advice. This work was funded by: NIH grant R21AI151725 to ALR; and CDC contract 200-2021-11195 to ALR.

## Conflict of Interest Statement

The authors declare no conflicts of interest.

**Supplementary Datafile 1**: The output BED files from a ViReMa analysis of three samples of SARS-CoV-2 previously described were used to illustrate the plotting features of ViReMaShiny. These files are also available at https://routhlab.shinyapps.io/ViReMaShiny and https://jayeung12.github.io

